# Identification of a non-exported Plasmepsin V substrate that functions in the parasitophorous vacuole of malaria parasites

**DOI:** 10.1101/2023.05.15.540838

**Authors:** Margarida Ressurreição, Aline Fréville, Christiaan van Ooij

**Author notes:** Contributed equally (Margarida Ressurreição, Aline Fréville).

## Abstract

Malaria parasites alter multiple properties of the host erythrocyte by exporting proteins into the host cell. Many exported proteins contain a five-amino acid motif called the Plasmodium Export Element (PEXEL) that is cleaved by the parasite protease Plasmepsin V (PM V). The presence of a PEXEL is considered a signature of protein export and has been used to identify a large number of exported proteins. The export of proteins becomes essential midway through the intraerythrocytic cycle – preventing protein export blocks parasite development 18-24 h after invasion. However, a genetic investigation revealed that the absence of the PEXEL protein PFA0210c causes parasite development to arrest immediately after invasion. We now show that this protein (renamed PV6) is cleaved by PM V but not exported into the host erythrocyte and instead functions in the parasitophorous vacuole. Furthermore, we show that the lysine residue that becomes the N terminus of PV6 after processing by PM V prevents export. This is the first example of a native *Plasmodium falciparum* PM V substrate that remains in the parasitophorous vacuole. We also provide evidence that the parasite produces at least one additional essential, non-exported PM V substrate. These results reveal that the presence of a PEXEL, and hence processing of a protein by PM V, does not always target a protein for export and that PM V has a broader function in parasite growth beyond processing exported proteins. Furthermore, we utilized this finding to investigate possible requirements for protein export further.

## INTRODUCTION

Export of proteins from malaria parasites into the host erythrocyte is an essential aspect of the parasite-host interaction – exported proteins induce changes in the permeability of the host cell, adhesiveness to the endothelium and deformability of the cell (Maier et al., 2009; Spillman et al., 2015). Exported proteins are synthesized in the parasite and transported through the ER and secretory pathway, released into the parasitophorous vacuole and subsequently transported past the parasitophorous vacuole membrane (PVM) through a large protein complex, the Parasite Translocon of EXported proteins (PTEX), into the host erythrocyte cytosol (Koning-Ward et al., 2009; Beck et al., 2014; Elsworth et al., 2014; Ho et al., 2018; Beck and Ho, 2021). The PTEX consists of three essential core subunits, HSP101, PTEX150 and EXP2, with several accessory proteins, of which PTEX88 and Trx2 are the most prominent (Koning-Ward et al., 2009; Matz et al., 2013; Batinovic et al., 2017; Chisholm et al., 2018; Ho et al., 2018). An important step in our understanding of protein export was the discovery of a motif shared by many exported proteins, the *Plasmodium* Export Element (PEXEL; also referred to as Host Targeting (HT) motif) (Hiller et al., 2004; Marti et al., 2004). The PEXEL is a five-amino acid motif with the sequence RxLxE/D/Q (a K is sometimes allowed at the first position (Schulze et al., 2015)). The discovery of the PEXEL allowed the prediction of the set of exported proteins produced by malaria parasites, which revealed that these parasites encode hundreds of exported proteins (Hiller et al., 2004; Marti et al., 2004; Sargeant et al., 2006; Boddey et al., 2013). The PEXEL is cleaved in the ER by the parasite protease Plasmepsin V (PM V) (Chang et al., 2008; Boddey et al., 2010; Russo et al., 2010; Sleebs et al., 2014; Tarr and Osborne, 2015) during translocation (Sleebs et al., 2014; Marapana et al., 2018). It has been speculated that cleavage by PM V designates proteins for export (Boddey et al., 2010; Russo et al., 2010) and to date, no substrates of PM V other than exported proteins have been identified. However, inhibition of PM V during the schizont stage leads to a different phenotype than inhibition of protein export – when PM V is inhibited starting at an early schizont stage, parasite development arrests immediately after invasion of the parasite (Polino et al., 2020), in contrast to the phenotype of inhibition of the PTEX during the schizont stage, which leads to an arrest late in the ring stage (Beck et al., 2014; Elsworth et al., 2014; Charnaud et al., 2018; Garten et al., 2018). This indicates that PM V may perform functions in addition to preparing proteins for export.

In addition to the PEXEL proteins, malaria parasites encode a second type of exported protein: the PEXEL-Negative Exported Proteins (PNEPs) (Haase et al., 2009; Spielmann and Gilberger, 2010; Heiber et al., 2013). Soluble PNEPs may be targeted to the ER by canonical N-terminal signal sequences but many PNEPs lack a signal sequence and instead contain an internal hydrophobic sequence that targets the protein to the ER. After transport through the secretory pathway and delivery to the PV, PNEPs are also translocated through the PTEX (Elsworth et al., 2014).

Biochemical and structural investigations of the PTEX have provided important insight into the mechanism of protein transport (Ho et al., 2018; Beck and Ho, 2021). Proteins require unfolding (Knuepfer et al., 2005a; Gehde et al., 2009; Grüring et al., 2012), mediated by HSP101 (Ho et al., 2018; Matthews et al., 2019), which also provides the energy for the translocation of the peptide chain through the pore of the PTEX complex, which is formed by EXP2 (Mesén-Ramírez et al., 2016; Ho et al., 2018). Exactly how parasite proteins are recognized by the PTEX has remained unclear. It has been speculated that processing by PM V acts as a signal that targets the protein for export (Russo et al., 2010; Gabriela et al., 2022) and that exported proteins are potentially already recognized by HSP101 in the ER (Gabriela et al., 2022). Several studies have indicated that an important factor that determines export is the nature of the N-terminal residue of the protein (Grüring et al., 2012; Tarr et al., 2013). Processing of the protein by PM V in proteins that contain a PEXEL, which occurs between the L and the second x residue in the PEXEL (Boddey et al., 2010; Russo et al., 2010), produces a protein in which x becomes the new N-terminal residue; changing the protease that processes model exported proteins had no effect on the export of the protein if the new N-terminal residue in the processed protein remained the same (Tarr et al., 2013). However, these experiments were performed using native exported proteins. In this study we show genetically, biochemically and microscopically that PFA0210c is a native *Plasmodium falciparum* PM V substrate that is not exported but instead functions in the PV and provide evidence that the parasite produces at least one other PM V substrate that is not exported. Our findings show that processing of substrates by PM V does not per se target a protein for export and that the protease has an additional role in processing parasite proteins important for the development of the parasite immediately after invasion. Furthermore, we use our and previous findings to analyze the sequences of exported and non-exported proteins to refine the signal for protein export further.

## MATERIALS AND METHODS

### Parasites

All experiments were performed with *P. falciparum* strain 3D7 and the transfected parasite line PfBLD529 (PV6-diCre) containing the floxed *pfa0210c* gene described previously (Hill et al., 2016). Parasites were maintained as described previously (Ressurreição et al., 2020) in RPMI-1640 medium (Life Technologies) supplemented with 2.3 g/L sodium bicarbonate, 4 g/L dextrose, 5.957 g/L HEPES, 0.5% AlbuMax II, 50 μM hypoxanthine and 2 mM L-glutamine (cRPMI). Cultures were maintained at 3% hematocrit and incubated at 37°C. Human erythrocytes were obtained from the National Blood Transfusion Service, UK and from Cambridge Bioscience, UK.

Parasites were synchronized by isolating schizonts on a Percoll gradient and allowing these parasites to invade uninfected erythrocytes for 1 to 3 hours. Remaining schizonts were removed on a second Percoll gradient. Rapamycin treatment was performed as described previously (Collins et al., 2013a; Knuepfer et al., 2017). Briefly, rapamycin was added to parasite cultures to a final concentration of 10 nM (from a stock solution of 100 µM in DMSO) and cultures were incubated at 37°C for one hour. Parasites were then pelleted and resuspended in cRPMI. Control cultures were treated similarly using DMSO instead of rapamycin.

Parasites were transfected using the schizont transfection protocol (Collins et al., 2013b). Briefly, parasites were synchronized using a Percoll gradient purification to set up invasion of fresh erythrocytes for 1-3 h followed by sorbitol lysis of remaining schizonts. To prepare parasites for transfection, schizonts were purified on a Percoll gradient and allowed to invade fresh erythrocytes. After 1-3 h, the remaining schizonts were removed from the culture on a Percoll gradient and transferred to a 15 ml tube. These schizonts were washed with cRPMI, transferred to microcentrifuge tubes and pelleted. The schizonts were subsequently resuspended in 100 µl AMAXA nucleofection P3 Primary Cell transfection reagent and 10 µl TE containing 15-30 µg targeting plasmid and 20 µg plasmid pDC2-Cas9-hDHFRyFCU that encodes the appropriate guide RNA and Cas9 (Knuepfer et al., 2017), followed by transfection using the nucleofection device (AMAXA), program FP 158. Transfected schizonts were mixed with uninfected erythrocytes and incubated with shaking at 37°C for 30 minutes to allow invasion. Drug selection using 2.5 nM WR99210 was initiated after 24 h and maintained for 7 days. Transfected parasites were in most cases recovered after 3 to 4 weeks.

### Plasmids

Plasmids pBLD634 encoding full-length PV6 with a K64A mutation, controlled by the native promoter, was made as follows. First, the K-A mutation was introduced using overlapping PCR. The 5’ and 3’ regions were amplified using primer pairs CVO081-CVO520 and CVO519-CVO023 (see Supp Table 1 for primer sequences), respectively, using *P. falciparum* 3D7 gDNA and a PV6-HA_3_-His6-GFP fusion (van Ooij et al., 2013) as templates. The resulting fragments were cloned into pBLD619 (this plasmid was made by cloning 1938 bp upstream of *pv6* into pBLD466 (also described as pBSPfs47DiCre (Knuepfer et al., 2017)), which contains two homology regions for integration of the plasmid into the *pfs47* locus (Knuepfer et al., 2017)) to produce pBLD634. These plasmids were integrated into the *pfs47* locus as described, using plasmid pDC287, which encodes Cas9 and a guide RNA targeting *pfs47*, as described previously (Knuepfer et al., 2017).

EXP1-HA_3_-PV6, AMA1-HA_3_-PV6 and PV6-HA_3_-PV6 expression plasmids were made as follows. Sequences encoding the EXP1-HA_3_-PV6 and the 5’ region of PV6 (spanning the fraction of the gene from the start ATG to the PEXEL) were synthesized as a gene fragment codon optimized for *Escherichia coli* (with further manual curation) to decrease similarity with the native *exp1* and *pv6* genes. The fragments were cloned into pCR-Blunt (Invitrogen), producing pBLD701 and pBLD703, respectively. Subsequently the 1086 bp downstream of the *pv6* gene was amplified using primers CVO675 and CVO676 and introduced into pBLD701 at the AflII site using InFusion (Takara), producing pBLD702. The fragment containing EXP1-HA_3_-PV6 and sequence downstream of *pv6* was released and cloned into pBLD619, which contains two homology regions for integration of the plasmid into the *pfs47* locus (Knuepfer et al., 2017) at the AvrII and AflII sites, producing pBLD706. Subsequently, the region encoding the EXP1 signal sequence was replaced by the recodonized region encoding the PV6 N-terminus (from pBDL703) or the AMA1 N-terminus using AvrII and XhoI (amplified from gDNA using primers CVO714 and CVO730), yielding pBLD708 and pBLD728, respectively. pBLD708 and pBLD728 were integrated into the *pv6* locus using Cas9-mediated gene modification as described previously (Hill et al., 2016).

### PCR

Verification of modification of the *pv6* locus was performed by first isolating genomic DNA from transfected parasites (Monarch genomic DNA isolation kit, New England Biolabs). Integration of targeting plasmid at the *pfs47* locus was determined with primer pairs CVO119-CVO120 (wildtype locus and integrated locus), CVO119-CVO070 (integration-specific) and CVO120-CVO694 (integration-specific). PCRs were set up according to manufacturers’ instructions, using Q5 Polymerase (New England Biolabs). Integration of targeting plasmid at the *pv6* locus was tested with the primer pairs CVO689-CVO111 (wildtype 5’ end), CVO071-CVO183 (wildtype 3’ end), CVO689-CVO586 (integrant 5’ end) and CVO321-CVO183 (integrant 3’ end) using Q5 Polymerase (New England Biolabs). PCRs were set up according to manufacturers’ instructions, using Q5 Polymerase (New England Biolabs) to amplify the 3’ end and CloneAmp (Takara Bio) for amplification of the 5’ end.

### Microscopy

Images of Giemsa-stained smears were captured using an Olympus BX51 microscope equipped with a 100x oil objective and an Olympus SC30 camera controlled by CellSense software.

For colocalization of PV6 with apical organelles markers, late-stage parasites were purified on a Percoll gradient. The parasites were smeared on a microscope slide, air-dried and stored at −20°C. Parasites were fixed with 4% paraformaldehyde in PBS, permeabilized with 0.1% Triton X-100, blocked with 3% BSA and subsequently stained with antibodies against PV6 and an apical organelle marker (AMA1 (dilution 1:500), RAP1 (1:100), Ron4 (1:500), all three kind gifts of Michael Blackman (Francis Crick Institute) and RESA (1:500), obtained from the Antibody Facility at The Walter and Eliza Hall Institute of Medical Research)), followed by staining with fluorescently-labeled secondary antibodies and staining of the DNA with Hoechst 33342 (5 µg/ml). To visualize PV6 after invasion, purified schizonts were allowed to invade fresh erythrocytes for approximately 1 hour and subsequently fixed in solution with 4% paraformaldehyde and 0.01% glutaraldehyde in PBS for one hour at room temperature (Tonkin et al., 2004). The parasites were permeabilized and stained as described above using anti-PV6 and anti-RESA or anti-EXP2 (1:2000) antibodies and Hoechst. The parasites were then stored at 4°C in a small amount of PBS (10-20 µl). For microscopy, 1.5 µl of parasites resuspended was placed on a polyethyleneimine-coated glass slide, mixed with 1.5 µl of Vectashield antifade mounting medium and covered with a cover glass, which was then sealed with nail polish. The parasites were imaged on a Nikon Ti-E inverted microscope with Hamamatsu ORCA-Flash 4.0 Camera and Piezo stage driven by NIS elements version 5.3 software. The images were processed using FIJI. The resulting images were cropped and file sizes were altered using Photoshop and figures were produced using Illustrator.

Export of PV6 K64A was visualized by fixing 3D7 parasites and parasites expressing PV6 K64A in solution and prepared for microscopy as described above for the visualization of PV6 after invasion. The samples were stained with a mouse monoclonal anti-PV6 antibody and rabbit anti-PV1 antiserum (a kind gift of Eizo Takashima) or affinity-purified rabbit anti-PV6 and mouse anti-EXP1 antibodies (a kind gift from Mike Blackman).

### Measurement of parasite size

To measure the size of parasites producing or lacking PV6 after invasion, PfBLD529 parasites were synchronized and treated with rapamycin or DMSO approximately 30 h after initiation of invasion. Egress of the parasites was inhibited with the addition of Compound 2 to 1.5 µM (Collins et al., 2013b). To initiate highly synchronized invasion, Compound 2 was removed by washing the cells with cRPMI, after which samples were taken at the indicated time points. One hour after the removal of compound 2, it was added to the samples once again to prevent further invasion. To prevent the repeated sampling from affecting parasite development, the DMSO-treated and rapamycin-treated culture were each split into four small flasks, which were sampled sequentially. At the indicated time points, parasites were smeared and stained with Giemsa. The diameter of the parasite was determined by imaging the Giemsa-stained parasites using an Olympus BX51 microscope equipped with an 100x oil objective and an Olympus SC30 camera controlled by CellSense software. The size was determined by measuring the longest diameter in each parasite using FIJI/ImageJ software, using the line selection tool and set measurement function.

### Growth assays

To determine parasite growth rates, parasites were synchronized using Percoll gradients as described above. Parasites containing a floxed version of the *pv6* gene were treated with rapamycin or DMSO as described above immediately after invasion. The cultures were diluted to a parasitemia of 0.1%-0.5% and growth was measured by removing a 50 µl aliquot and fixing that in an equal volume of 2x fixative and stain (8% paraformaldehyde, 0.2% glutaraldehyde, 2x SYBR Green). Fixed samples were stored at 4°C. To prepare the samples for cytometry, the fixative was removed and the cells were resuspended in 1 ml PBS. A 200 µl aliquot was transferred to a 96-well plate and the parasitemia was measured using an Attune cytometer outfitted with a CytKick autosampler (ThermoFisher). Laser settings used for detection of the erythrocytes and parasites were as follows: forward scatter 125 V, side scatter 350 V, blue laser (BL1) 530:30 280V. The experiment was set up in triplicate, with a minimum of three biological replicates.

### Streptolysin O and saponin treatment and immunoblots

SLO (SigmaAldrich) was titrated to quantify the hemolytic units per µl. To perforate the erythrocyte plasma membrane to release erythrocyte cytosol proteins, 10^8^ parasites were pelleted and resuspended in 100 µl PBS containing 4 hemolytic units SLO. To release erythrocyte cytosol and PV proteins, 10^8^ parasites were pelleted and resuspended in 100 µl 0.1% saponin in PBS. As a control, 10^8^ parasites were pelleted and resuspended in 100 µl PBS. The treated cells were pelleted; the supernatant of these samples was collected and mixed with an equal volume 2X SDS-PAGE loading dye and the pellet was resuspended in 1X SDS-PAGE loading dye.

### Immunoblots

Parasite lysates were separated by SDS-PAGE, transferred to nitrocellulose and probed with anti-PV6 (1:1000), anti-HRPII (Immunology Consultants Laboratory, 1:2000), anti-aldolase-HRP (AbCam, 1:5000), anti-PV1 (1:1000) or anti-EXP2 (1:2000) antibodies. Proteins were visualized by incubating the blots with the appropriate HRP-linked anti-mouse or anti-rabbit secondary antibodies and developing using Clarity ECL Western blotting substrates (Bio Rad).

### Plasmepsin V inhibitor treatment

To determine whether PV6 is a PM V substrate, infected erythrocytes in a synchronized parasite culture were isolated on a Percoll gradient. Approximately 30 hours after invasion, parasites were pelleted and either resuspended in 1X SDS-PAGE loading dye or resuspend in cRPMI containing 0, 20 or 50 µM PM V inhibitor WEHI-1252601 (a kind gift of Justin Boddey). These parasites were incubated at 37°C for 3 hours and then pelleted and resuspended in 1x SDS-PAGE loading dye. Lysates were separated by SDS-PAGE, transferred to nitrocellulose and probed with anti-PV6 (1:1000) or anti-EXP2 antibodies (1:2000).

To determine the effect of PM V inhibition on invading parasites, cultures of 3D7 and AMA1-HA_3_-PV6 parasites were synchronized and half the culture was treated with 7.5 µM WEHI-1252601 approximately 40 h after invasion. Egress was blocked by the addition of ML10 to 25 nM (Baker et al., 2017; Ressurreição et al., 2020); when most parasites appeared to have arrested, the parasites were pelleted and ML10 and WEHI-1252601 were removed. The culture that had been treated only with ML10 was split into two and WEHI-1252601 was added to one of the two cultures. The parasites were incubated at 37°C to allow egress and invasion of fresh erythrocytes. Parasites were collected one hour after the removal of ML10, smeared on slides and stained with Giemsa.

To determine the effect of inhibition of PM V on parasites expressing or lacking PV6, similar experiments were carried on PfBLD529 parasites. The parasites were synchronized and treated with rapamycin or DMSO approximately 30 hours after initiation of invasion and allowed to develop to schizont stage, at which point they were subjected to ML10 blocking and WEHI-1252601 treatment for 4 hours at 37°C. After removal of ML10 and WEHI-1252601, the parasites were allowed to egress and invade fresh erythrocytes for one hour. The Giemsa-stained parasites were imaged and the measurement of parasite size was performed as described above.

## RESULTS

### A *pfa0210c* mutant does not develop after invasion

In a previous investigation of a *pfa210c* mutant, we observed that its phenotype became apparent during the ring stage of the intraerythrocytic developmental cycle, very soon after invasion of the parasite (Hill et al., 2016). In contrast, mutants lacking subunits of the PTEX that are unable to export proteins develop seemingly as normal through the ring stage until the ring-trophozoite transition (18-24 h post-invasion), indicating that the functions of exported proteins are not essential until this stage (Beck et al., 2014; Elsworth et al., 2014; Sleebs et al., 2014; Charnaud et al., 2018). Hence, despite being annotated as an exported PEXEL protein, this phenotype of the *pfa0210c* mutant implies that PFA0210c performs an essential function prior to the time at which protein export is required. We followed the development of the mutant parasites, starting from invasion of the merozoite, to gain a more detailed understanding of the timing of the appearance of this phenotype and determine whether the parasites undergo a temporary delay in development or suffer a terminal block in development. We synchronized parasites in which the *pfa0210c* gene could be deleted by the addition of rapamycin (Collins et al., 2013a; Hill et al., 2016), induced deletion of the *pfa0210c* gene approximately 30 hours after invasion and subsequently synchronized parasite egress using the cGMP-dependent protein kinase (PKG) inhibitor compound 2 (Collins et al., 2013b). No difference was detected between rapamycin-treated and DMSO-treated schizonts by Giemsa staining (Fig 1a), indicating that the loss of the *pfa0210c* gene does not affect the development of late-stage parasites and merozoites. After removal of compound 2, the parasites displayed a phenotype within one hour (as invasion commences approximately 20 minutes after the removal of compound 2, the parasites in this sample had invaded within the preceding 40 minutes). No mutant parasites were detected that resembled wild-type parasites (Fig 1A). Measurement of the parasites confirmed the difference between the mutant and wildtype parasites; the size of the mutant parasites did not change during the remainder of the period during which the parasites were followed, indicating that the phenotype is terminal rather than a delay in development (Fig 1B), consistent with the finding that parasites lacking PFA0210c do not grow (Hill et al., 2016). Therefore, we conclude that PFA0210c functions prior to the requirement for protein export.

**Figure 1.**
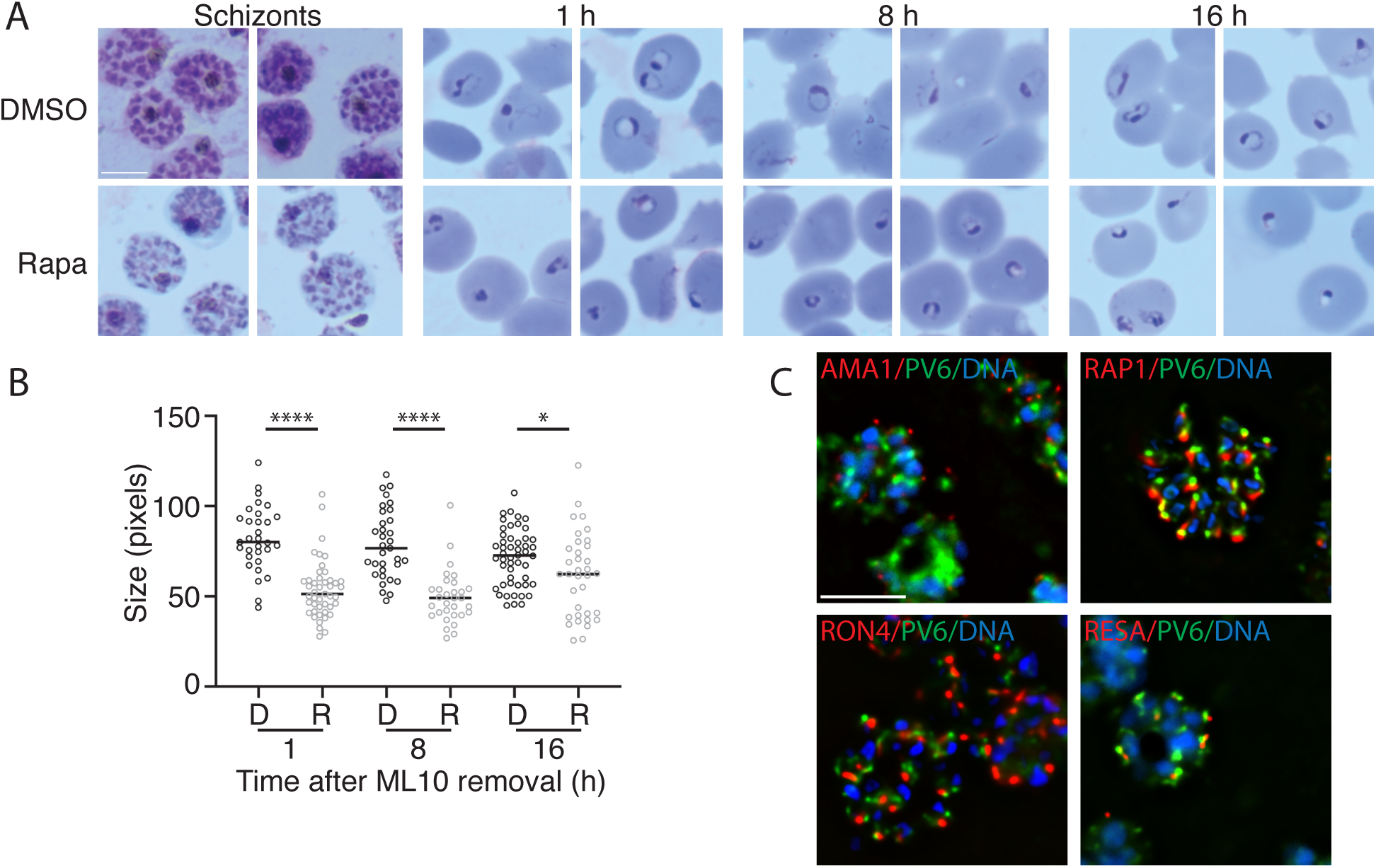
Development of parasites expressing or lacking PV6 during ring stage and PV6 localization in schizonts. A) Synchronized parasites with a floxed *pv6* (*pfa0210)* gene were treated at the late trophozoite stage with DMSO or with rapamycin to induce excision of *pv6*. Parasites were examined at the schizont stage in Giemsa-stained smears and subsequent egress was synchronized with ML10 treatment. Giemsa smears were produced 1, 8 and 16 hours after ML10 removal. Scale bar represents 5 µm. B) Quantitation of the size of the parasite diameter in the samples shown in panel A. Data are based on the measurement of at least 30 rings. Mann–Whitney U test was performed for statistical analysis: *p < 0.05, ****p < 0.0001. D-DMSO treated, R-Rapamycin treated. C) Costaining of schizonts with antibodies against PV6 (green) and the apical organelles markers AMA1 (micronemes), RAP1 (rhoptry), RON4(Rhoptry neck) and RESA (dense granule) (all in red) by immunofluorescence microscopy. DNA was stained with Hoechst 33342 (blue). Scale bar represents 5 µm.

As PFA0210c functions very soon after invasion, it is very likely present in merozoites at the time of invasion. We therefore carried out colocalization studies using markers of apical organelles to determine its localization. This revealed that the protein is indeed present in merozoites, but we were not able to detect consistent colocalization of PFA0210c with the standard markers of apical organelle; no colocalization was detected with markers of the microneme (AMA1), rhoptry (RAP1), rhoptry neck (RON4) or dense granules (RESA) (Fig 1C). As RESA and PV6 appeared to be closely positioned together, we determined whether PV6 colocalizes with EXP2, another dense granule markers. This also did not colocalize with PV6, even when localization was determined in merozoites (Supp Fig 1).

### PFA0210 is cleaved by Plasmepsin V but not exported

PFA0210c was initially identified as a PEXEL protein and subsequent studies have identified it as a substrate of PM V using mass spectrometry (Silvestrini et al., 2010; Shears et al., 2019). However, as the phenotype of the *pfa0210c* mutant indicates that the protein does not function as an exported protein, unlike other known PM V substrates, we aimed to verify biochemically that PFA0210c is indeed cleaved by PM V. For this we treated parasites with the PM V inhibitor WEHI-1252601 (601) for three hours, starting approximately 30 hours after invasion; at this time point PFA0210c is already produced (van Ooij et al., 2013) and protein export has commenced, decreasing the chance that addition of the inhibitor has a detrimental effect on the parasites. However, some PFA0210c that has already been processed will be present in these samples (Fig 2A, upper panel, time 0). Immunoblotting revealed the appearance of a band of with the expected molecular mass of full-length, uncleaved PFA0210c (53.6 kD) in the extracts of parasites treated with the inhibitor (Fig 2A, upper panel); no size shift was detected for EXP2, a protein that is cleaved by signal peptidase, in these samples (Fig 2A, lower panel). These results confirm that PFA0210c is indeed a PM V target.

**Figure 2.**
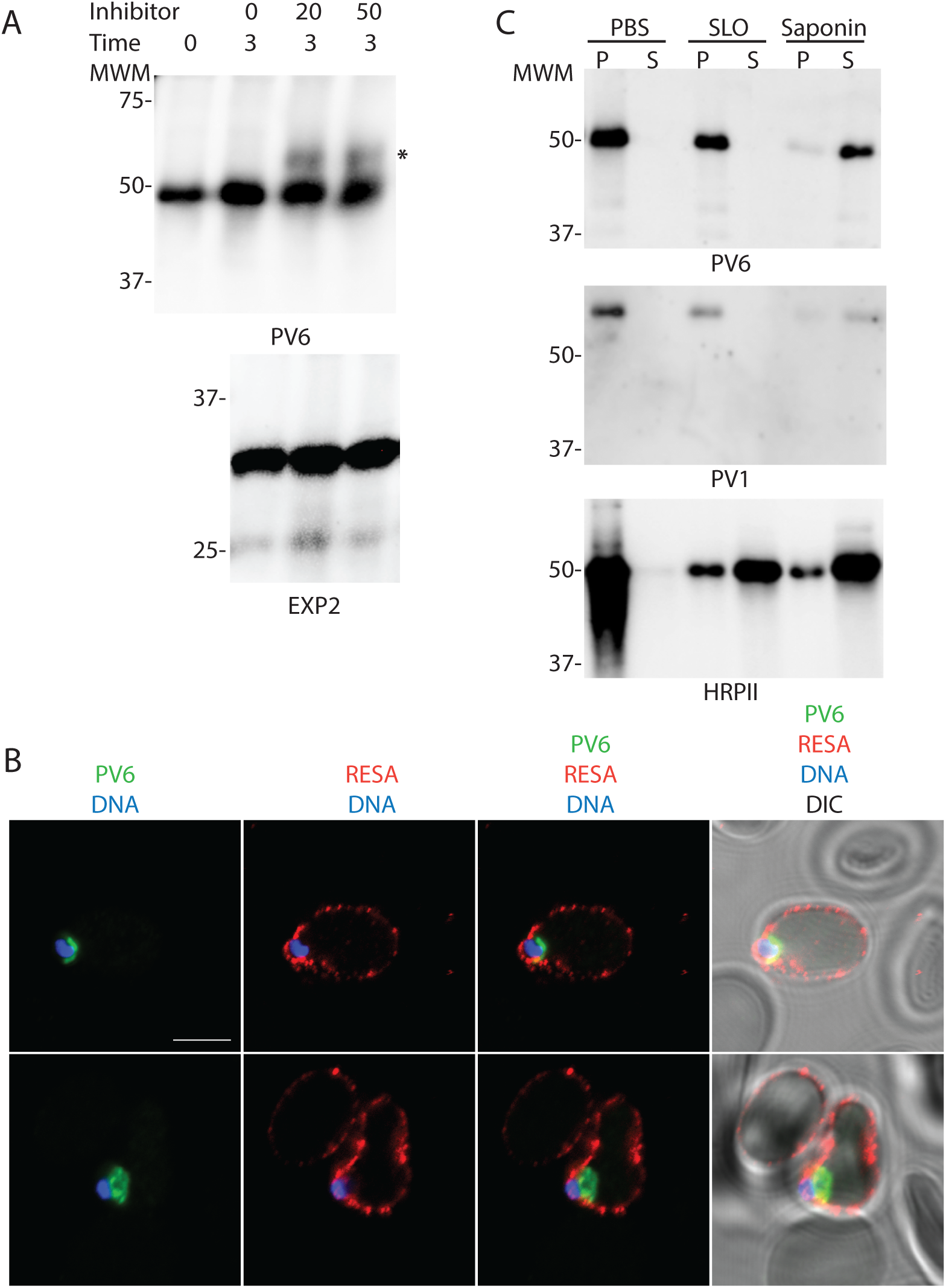
Role of Plasmepsin V in processing of PV6 and localization of PV6 after invasion. A) Parasites were treated with the indicated amount (in µM) of the PM V inhibitor WEHI-1252601 for three hours. Parasite extracts were separated by SDS-PAGE and probed with antibodies against PV6 (top panel) and EXP2 (bottom panel). Note the presence of a higher molecular mass band in the inhibitor-treated samples in the anti-PV6 (indicated with *) but not the anti-EXP2 immunoblot. Expected sizes of unprocessed PV6 and PV6 processed by PM V: 53.6 kD and 46.4kD, respectively. B) Localization of PV6 and RESA immediately after invasion of 3D7 parasites. Invasion was synchronized with ML10 and samples were collected within two hours after removal of ML10. Note that the erythrocyte at the top of the panel is also infected and therefore also displays anti-RESA staining, but that the parasite in that cell is outside of the plane of focus (see Supp Fig 2A for a maximum projection of the image). Scale bar represents 5 µm. C) Differential lysis of infected erythrocytes with Streptolysin O and saponin. Infected erythrocytes were treated with PBS, SLO or saponin, the erythrocytes were pelleted and the supernatant and pellet were collected. Samples were prepared for immunoblotting and probed with antibodies against the indicated proteins. Note the similarity in the release of PV6 and the parasitophorous vacuole marker PV1 in the samples.

We next determined the localization of PFA0210c after invasion using immunofluorescence. This revealed that PFA0210c remains in the PV (Fig 2B) even in cells where the exported protein RESA has been exported and has bound to the host cell cytoskeleton, with no detectable RESA remaining in the PV (Fig 2C, Supp Fig 2A); no PFA0210c was detected in the erythrocyte. The localization of PFA0210c was similar to that of EXP2, although interestingly, little to no overlap of the staining was detected (Fig 2B), indicating that PV6 may be localized outside of the PV/PVM domains containing EXP2 previously identified (Garten et al., 2020).

However, it is often difficult to visualize soluble exported proteins in the cytoplasm of the host cell, especially at the early stages of the intraerythrocytic cycle when soluble proteins will be greatly diluted in the host cell cytoplasm compared to the PV. We therefore determined biochemically whether the protein is retained in the PV using selective permeabilization of the host cell and PVM with Streptolysin O (SLO) and saponin, respectively (Ansorge et al., 1996, 1997; Jackson et al., 2007). Treatment of infected parasites with PBS, which leaves the erythrocyte intact, or SLO, which perforates the erythrocyte membrane, and thereby releases exported proteins into the supernatant while leaving the PVM intact, did not release PFA0210c, even though the soluble exported protein HRPII was released (Fig 2C). In contrast, treatment of the parasite with saponin, which lyses the erythrocyte membrane and the PVM, released PFA0210c, along with the previously characterized PV protein PV1 (Chu et al., 2011; Morita et al., 2018). This further indicates that PFA0210c is a PV-resident protein. Collectively, these results show that PFA0210c is a PM V target that remains in the PV and therefore we propose the name PV6 for this protein to reflect its subcellular localization.

### An AMA1-HA_3_-PV6 fusion can replace the function of native PV6

Previous studies revealed that, when overexpressed using a Cam promoter, a PV6-GFP fusion protein can be detected in the host cell cytosol (van Ooij et al, 2008). Although the results above show that PV6 is expressed and appears to function in the PV, we could not completely rule out potential additional functions of PV6 in the erythrocyte. We therefore produced a version of the gene that encodes a protein that is cleaved by signal peptidase, rather than PM V, to replace the native gene. For this, we fused the conserved region of PV6 starting from codon 122, including the START domain and the regulatory region in the C terminus but lacking the PEXEL to sequence encoding the AMA1 signal sequence (Fig 3A). In addition, we included sequence encoding a 3xHA tag (HA_3_) between the AMA1 signal sequence and the PV6 sequence, producing an AMA1-HA_3_-PV6 fusion (Fig 3A). As a control, we generated a fusion gene that contains the first 77 codons of PV6, including the PEXEL and 12 downstream codons, fused to sequence encoding the HA_3_ tag and PV6 starting from codon 122, the producing a PV6-HA_3_-PV6 fusion. These fusion genes were used to replace the native *pv6* gene using Cas9-mediated gene replacement, leading to rapid and efficient integration; no wildtype version of the gene could be detected by PCR in the transfected parasite populations (Fig 3B). The genetically modified parasites expressed the fusion proteins, as indicated by anti-HA immunoblots (Fig 3C). Of note, the modification of PV6 N-terminal region leads to the production of slightly smaller proteins, as displayed in the anti-PV6 blot (Fig 3C).

**Figure 3.**
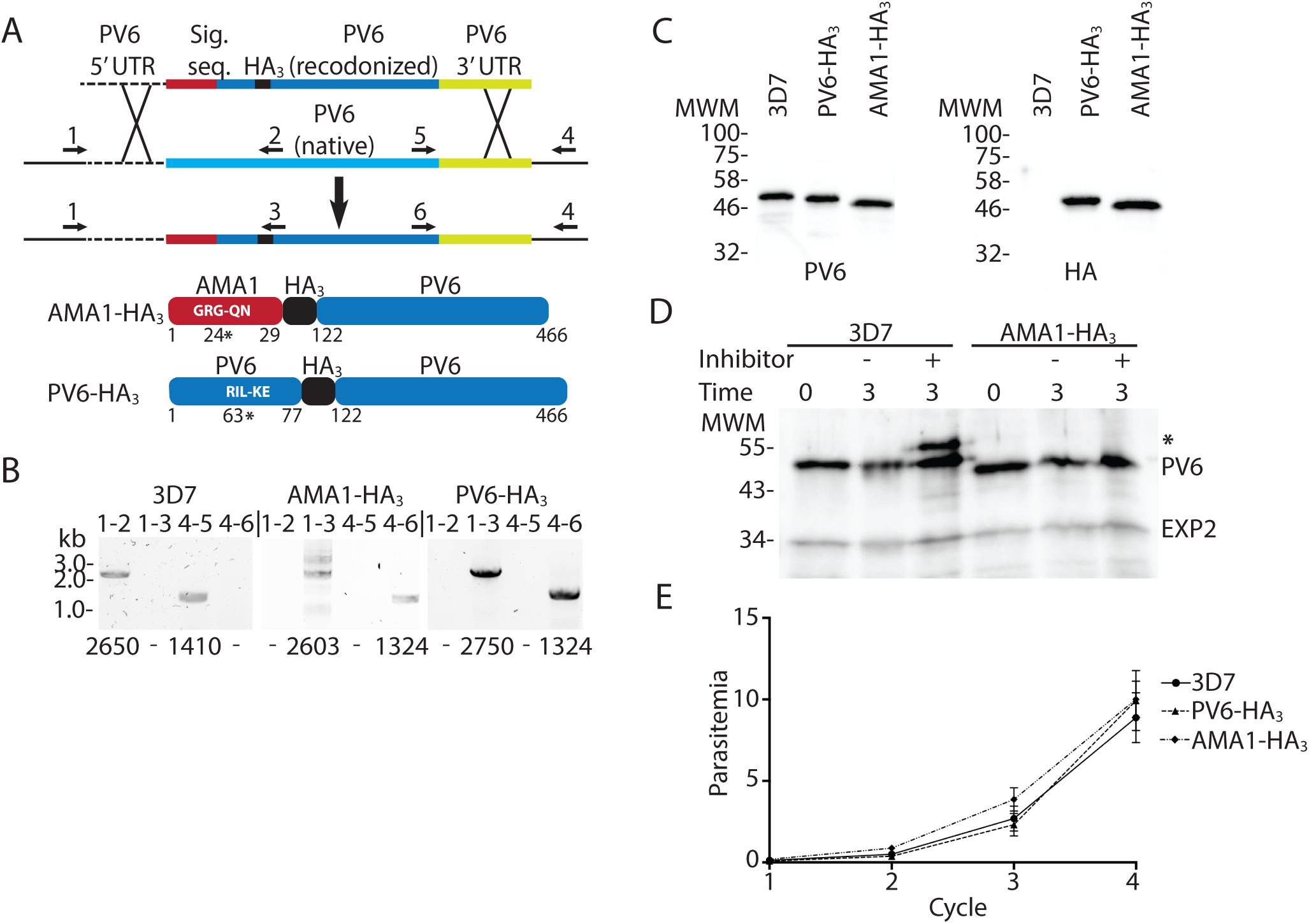
Replacement of native PV6 with an AMA1-HA_3_-PV6 fusion. A) Outline of the replacement strategy of the native *pv6* gene with the gene encoding the AMA1-HA_3_-PV6 fusion. The same strategy was used to create the PV6-HA_3_-PV6 fusion. Cartoons at the bottom show the expected protein products after replacement of the native gene with the inserted gene. Indicated in white letters is the predicted signal sequence cleavage site in the AMA1-HA_3_-PV6 fusion and the PEXEL sequence of the PV6-HA_3_-PV6 fusion. The number and * indicate the signal sequence cleavage site in the AMA1-HA_3_-PV6 and PM V cleavage site in the PV6-HA_3_-PV6 fusion. Numbers indicate amino acid residues of the native proteins. B) Integration PCR using wildtype and transfected parasite DNA. Primer pairs are indicated above the lanes (see also Supp Table 1), primer binding sites are indicated in panel A and expected sizes of PCR products are indicated below each lane. Note the absence of integration-specific PCR products in the samples using gDNA from wildtype parasites and the absence of wildtype-specific PCR products in the samples using gDNA from transfectants. C) Anti-PV6 (left) and anti-HA (right) immunoblots of wildtype (3D7) and transgenic parasite (PV6-HA_3_ and AMA1-HA_3_) extracts using the indicated antibodies. Expected molecular weights are 46.4 kDa (3D7 native PV6), 45.4 kDa (PV6-HA_3_-PV6) and 44.4 kDa (AMA1-HA_3_-PV6). D) Effect of Plasmepsin V inhibitor treatment on PV6, AMA1-HA_3_-PV6 and EXP2. Predicted sizes of uncleaved and cleaved proteins are 53.6 KDa and 46.4 kDa (PV6), 47.3 kDa and 44.4 kDa (AMA1-HA_3_-PV6) and 33.4 kDa and 30.8 kDa (EXP2). Note the absence of unprocessed AMA1-HA_3_-PV6 and EXP2 in the presence of the Plasmepsin V inhibitor. * indicates the higher molecular mass product in the 3D7 lysate. E) Growth assay of 3D7, PV6-HA_3_-PV6 and AMA1-HA_3_-PV6 parasites. Individual growth assays were set up in triplicate, data shown are from three biological replicates. Error bars indicate the standard error of mean but are in several cases too small to protrude from the symbol.

To ascertain that the AMA1-HA_3_-PV6 fusion is not cleaved by PM V, we treated wildtype parasites and parasites expressing the AMA1-HA_3_-PV6 fusion protein with the PM V inhibitor WEHI-1252601. Whereas a band of higher molecular mass was detected in extracts of wildtype PV6 parasites that had been treated with the inhibitor, no such band was detected in extracts of parasites expressing AMA1-HA_3_-PV6 (Fig 3D). Hence, AMA1-HA_3_-PV6 is not a PM V substrate.

As the genetically modified parasites were obtained quickly after the transfections (within 3-4 weeks) and without competing wildtype parasites, the growth rate of the parasites producing the fusion protein did not appear to be affected. Growth assays confirmed that the AMA1-HA_3_-PV6 and PV6-HA_3_-PV6 parasites do indeed grow at the same rate as the parent 3D7 parasite line (Figure 3E). Hence, the fusion protein that is targeted to the ER by a signal sequence, but is not cleaved by PMV, supports parasite growth. This indicates that cleavage by PM V is not essential for correct trafficking and function of the protein.

### Residue 4 of the PEXEL determines export of PV6

The PEXEL of PV6, RILKE, is unusual in that the fourth residue is a lysine. Although the PEXEL is usually denoted as RxLxE/Q/D (with a K allowed at the first position in some cases (Schulze et al., 2015)), previous studies have shown that the fourth residue affects the export of PEXEL proteins (Grüring et al., 2012; Tarr et al., 2013). Hence, we postulated that this residue may prevent export into the host cell. We therefore replaced the codon encoding the K residue with a codon encoding an A (K64A mutation), which is present at the fourth position of the PEXEL of many exported proteins, including the well-characterized exported protein KAHRP (Boddey et al., 2009) and introduced this mutated gene, controlled by the native *pv6* promoter, at the *pfs47* locus (Knuepfer et al., 2017), producing a merodiploid (Supp Fig 3A). By immunofluorescence microscopy, using rabbit anti-PV6 antibodies, PV6 was detected in the cytosol of erythrocytes infected with parasites that produce PV6 with the K64A mutation (Fig 4A, upper panel), in contrast to wildtype parasites, where the protein was detected only within the parasite and the PV, as identified by staining with the PVM marker EXP1 (Fig 4A, lower panel). We repeated the experiment with a monoclonal mouse anti-PV6 antibody, using PV1 as a marker for the PV, and once again we detected PV6 only in the parasite and the PV in untransfected parasites but detected PV6 also in the host cell cytosol in erythrocytes infected with parasites expressing the PV6 K64A mutant (Supp Fig 3B). These results indicate that the change in the N-terminal residue from K to A allows PV6 to be exported.

**Figure 4.**
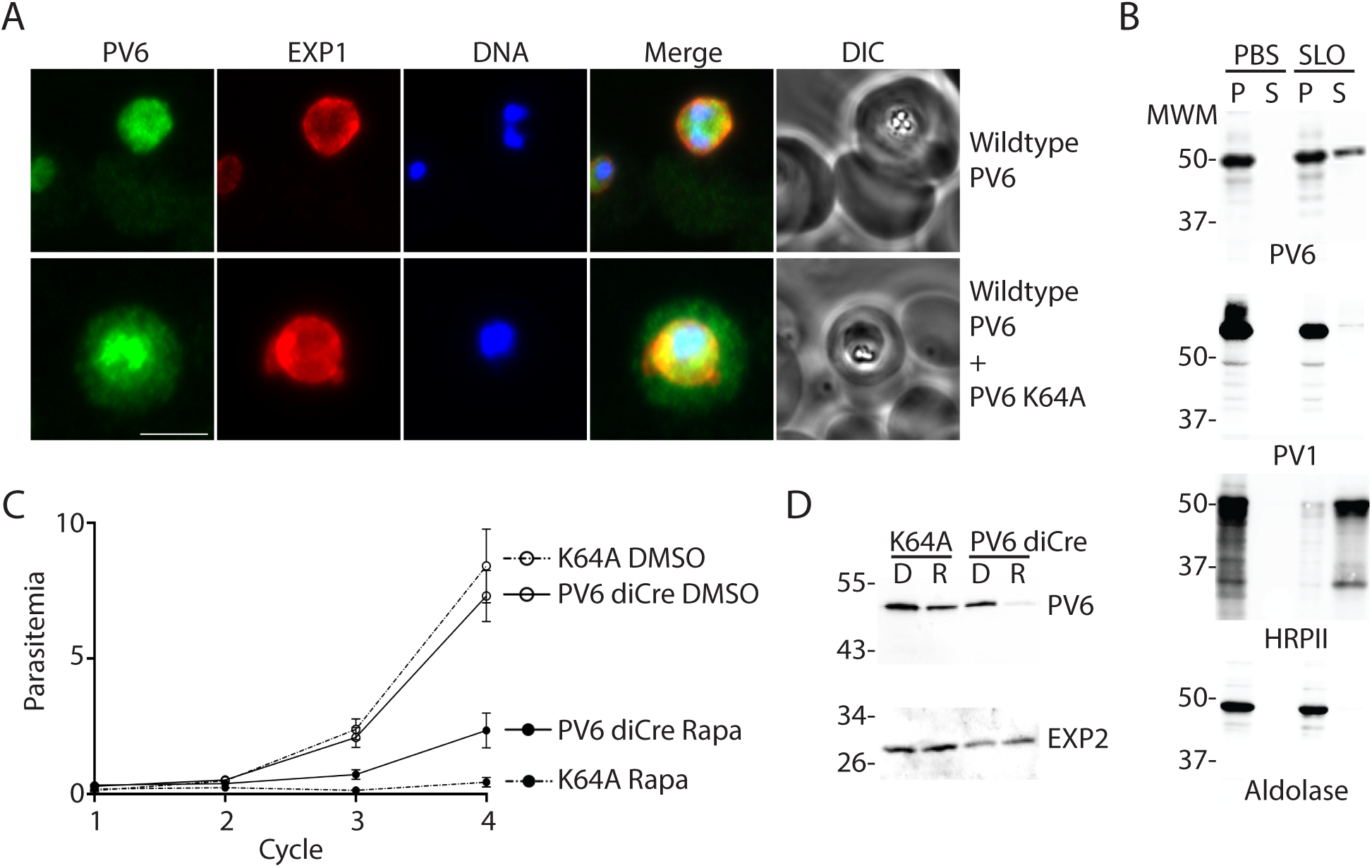
Localization and function of a PV6 K64A mutant. A) Immunofluorescence assays of parasites expressing wildtype PV6 or wildtype PV6 plus PV6 K64A. The PVM was visualized with an anti-EXP1 antibody. PV6 was detected in the erythrocyte cytosol only in erythrocytes infected with parasites expressing the PV6 K64A mutant. See Supp Fig 3 for additional images. Scale bar represents 5 µm. B) Selective permeabilization of erythrocytes infected with wildtype or parasites expressing the K64A mutant. Infected erythrocytes were treated with PBS or SLO and pelleted. The pellet and supernatant were subjected to SDS-PAGE and immunoblotted with the indicated antibodies. Note the release of PV6 in the supernatant of the permeabilized erythrocytes infected with parasites expressing PV6 K64A, whereas nearly all of the PV marker PV1 is retained in the parasite. C) Growth assays of PV6 diCre and parasites expressing PV6 K64A after treatment with DMSO or rapamycin to remove the native *pv6* gene. Growth assays were set up in triplicate; error bars indicate the error of mean but are in several cases too small to protrude from the symbol. D) Anti-PV6 (upper panel) and anti-EXP2 (lower panel) immunoblots of parental (PV6 diCre) and transfected parasite (K64A mutant line). Of note, PV6 expression is lost after rapamycin treatment in the parental line. When the same treatment is carried on the mutant line, the protein signal is only reduced reflecting the continued expression of the K64A mutant protein from the *pfs47* locus by the transgenic parasite.

To verify the microscopy results biochemically, we again performed differential lysis of infected erythrocytes using saponin and SLO. PV6 was readily detected in the supernatant of SLO-treated erythrocytes that were infected with parasites expressing the version of PV6 with the K64A mutation, indicating that the mutated version of the protein is exported (Fig 4B). As these parasites also express the native version of the protein, PV6 was also detected in the parasitophorous vacuole (Fig 4B, SLO pellet). As expected, the exported protein HRPII was also readily detected in SLO supernatant of both strains, whereas the PV protein PV1 protein was detected almost entirely in the pellet of the SLO-treated parasites of both strains (Fig 4B).

To test whether PV6 carrying the K64A mutation can support growth, we compared the growth rate of the K64A parasites with that of the parent parasite line, PfBLD529 PV6 diCre, in which the native *pv6* gene can be removed by treating the parasites with rapamycin. After we treated these parasites with DMSO or with rapamycin to remove the native *pv6* gene, no large difference between the two strains was observed in the growth of the parasites after treatment with either DMSO (the vehicle control) or after treatment of with rapamycin. Therefore, even though the K64A parasites continue to express the PV6 K64A mutant protein, this does not support growth of the parasites, indicating that the K64A PV6 mutant protein does not support growth (Fig 4C). As expected, immunoblot analysis confirmed the almost complete loss of PV6 expression in the parental line treated with rapamycin, whereas only a reduction is observed in the treated mutant (Fig 4D). There, the signal detected by the anti-PV6 antibodies corresponds most likely to the K64 PV6 protein encoded at the *pfs47* locus, as suggested by the diagnostic PCR analysis revealing a complete excision of the native *pv6* after rapamycin treatment of the K64A mutant (Supp Fig 3C).

Together, these results show that the native PV6 is not targeted for export, and hence retained in the PV, owing to the K residue at its N terminus.

### Analysis of the putative N termini of PEXEL proteins, PNEPs, PV proteins, PVM proteins and parasite surface proteins

The finding that the K residue at the N terminus of PV6 does not allow export of the protein prompted us to examine in more detail how often each amino acid is present at the fourth position of the PEXEL in proteins predicted to be exported and whether a K residue is present at the fourth position of known exported proteins. As has been noted previously, amino acids with small, hydrophilic side chains are common in this fourth position (Hiller et al., 2004; Marti et al., 2004; Boddey et al., 2009; Grüring et al., 2012; Tarr et al., 2013). Consistent with this, when we determined the frequency at which each amino acid is observed at the fourth position in predicted exported proteins in three different published exportomes (the Boddey, Sargeant and van Ooij exportomes), we found that most often a small, polar amino acid is found at this position (Hiller et al., 2004; Marti et al., 2004; Sargeant et al., 2006; Boddey et al., 2009). Overall, we found that ten different amino acids are observed at the fourth position of verified exported proteins (Table 1). Five residues, A, C, S, T and Y, are very common at this position, in part because they frequently occur in the large protein families STEVOR and RIFIN, although these residues are also present in many proteins for which export has been verified that are not part of large families, including KAHRP and PfEMP3 (A) (Knuepfer et al., 2005b; Boddey et al., 2009) and several FIKK kinases, MESA, REX3 and REX4 (S) (Coppel et al., 1988; Howard et al., 1988; Schulze et al., 2015; Davies et al., 2020). Export of proteins with other, less common, residues at position 4, including H (HRPII and HRPIII) (Akompong et al., 2002), G (RIFIN PFI0050c/PF3D7_0901000, PTP2 and the *Plasmodium vivax* protein PHIST CVC-8195/VX_093680) (Khattab and Klinkert, 2006; Akinyi et al., 2012; Koning-Ward et al., 2016; Wang et al., 2016), L (FIKK7.1) (Davies et al., 2020), N (GARP, FIKK9.7 and FIKK13) (Davies et al., 2016, 2020) and V (GEXP18) (Zhang et al., 2017) has also been observed. No known exported protein contains a large, charged residue at the fourth position (such as K), although each exportome predicts several such proteins. However, many of these predicted proteins have been shown or are predicted to be targeted to intracellular locations within the parasite or lack an annotated signal peptide or transmembrane domain and therefore are unlikely to be exported proteins (Supp Table 3). Five amino acids were not observed in the fourth position of the PEXEL proteins: D, M, P, Q and W. Based on this we categorized the amino acids into ‘Common-confirmed’ (A, C, S, T, Y), ‘Uncommon-confirmed’ (G, H, L, N, V), ‘Not confirmed – unlikely’ (E, F, K, R), ‘Not confirmed’ (I) and ‘Not observed’ (D, M, P, Q, W) (Table 1, also see Supp Table 4).

**Table 1.**
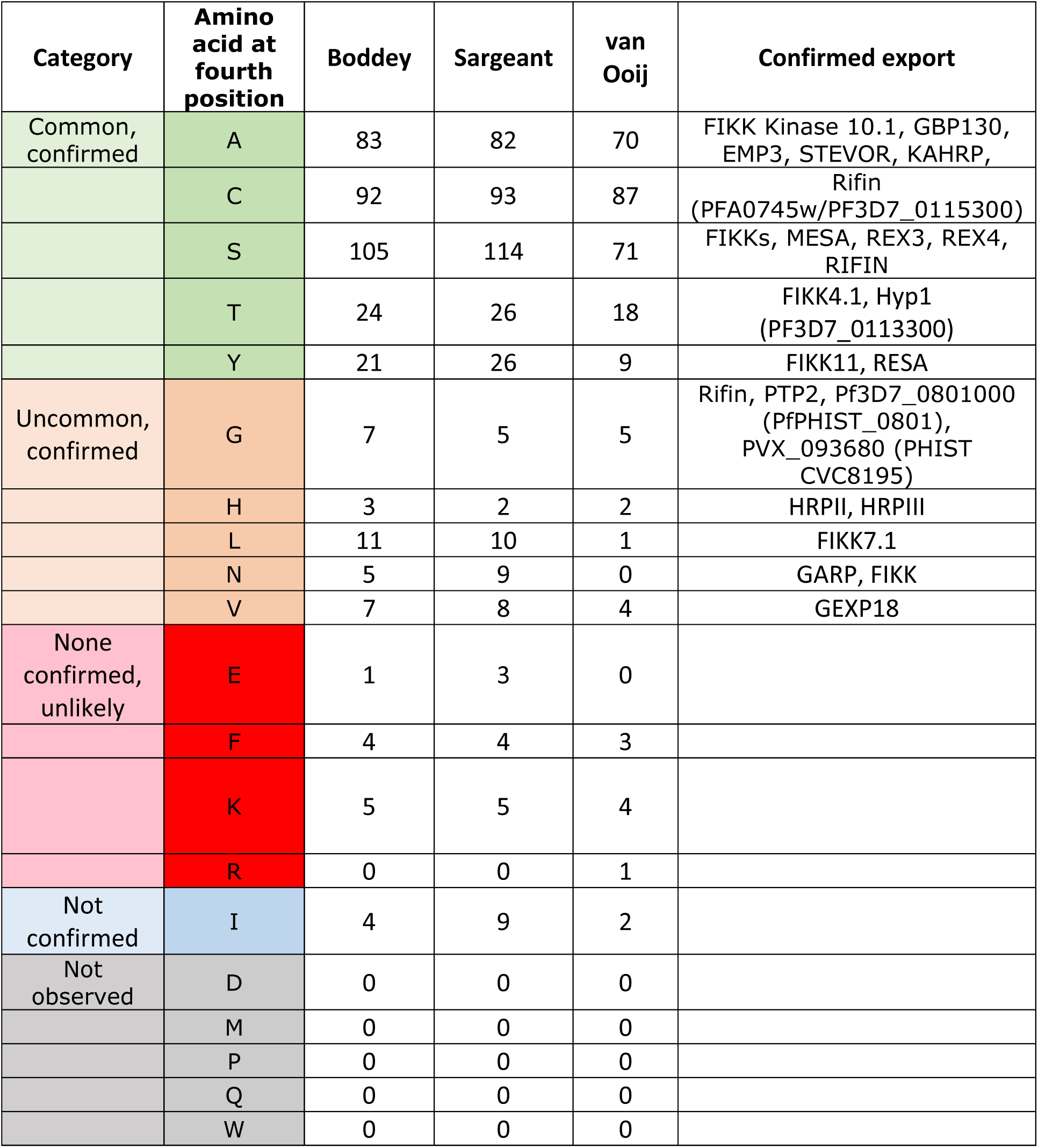
Occurrence of all amino acids in position 4 of the PEXEL in proteins predicted to be exported in the indicated exportomes. For references for conformation of protein export, please see text.

As PEXEL-Negative Exported Proteins (PNEPs) are also exported through the PTEX (Elsworth et al., 2014), we furthermore examined the N termini of several PNEPs and the members of the SURFIN family (Table 2). As the N-terminal methionine residue is removed by methionine aminopeptidase when it is followed by a small residue (primarily G, A, S, T, C, P and V) (Bonissone et al., 2013), we focused on the second residue of the protein. We found that most PNEPs contain a small amino acid after the start methionine and hence are likely to be processed by methionine aminopeptidase. Two PNEPs (PF3D7_0830500/PF08_0003 and PF3D7_1148900/PF11_0505) contain a residue following the M residue that has not been detected at position 4 of the PEXEL of exported proteins and that is not thought to be removed (N and E), making it likely that these proteins retain their start M. Although export was detected for these proteins, the proportion of the protein that was exported was much lower than that of the other PNEPs (Heiber et al., 2013).

**Table 2.**
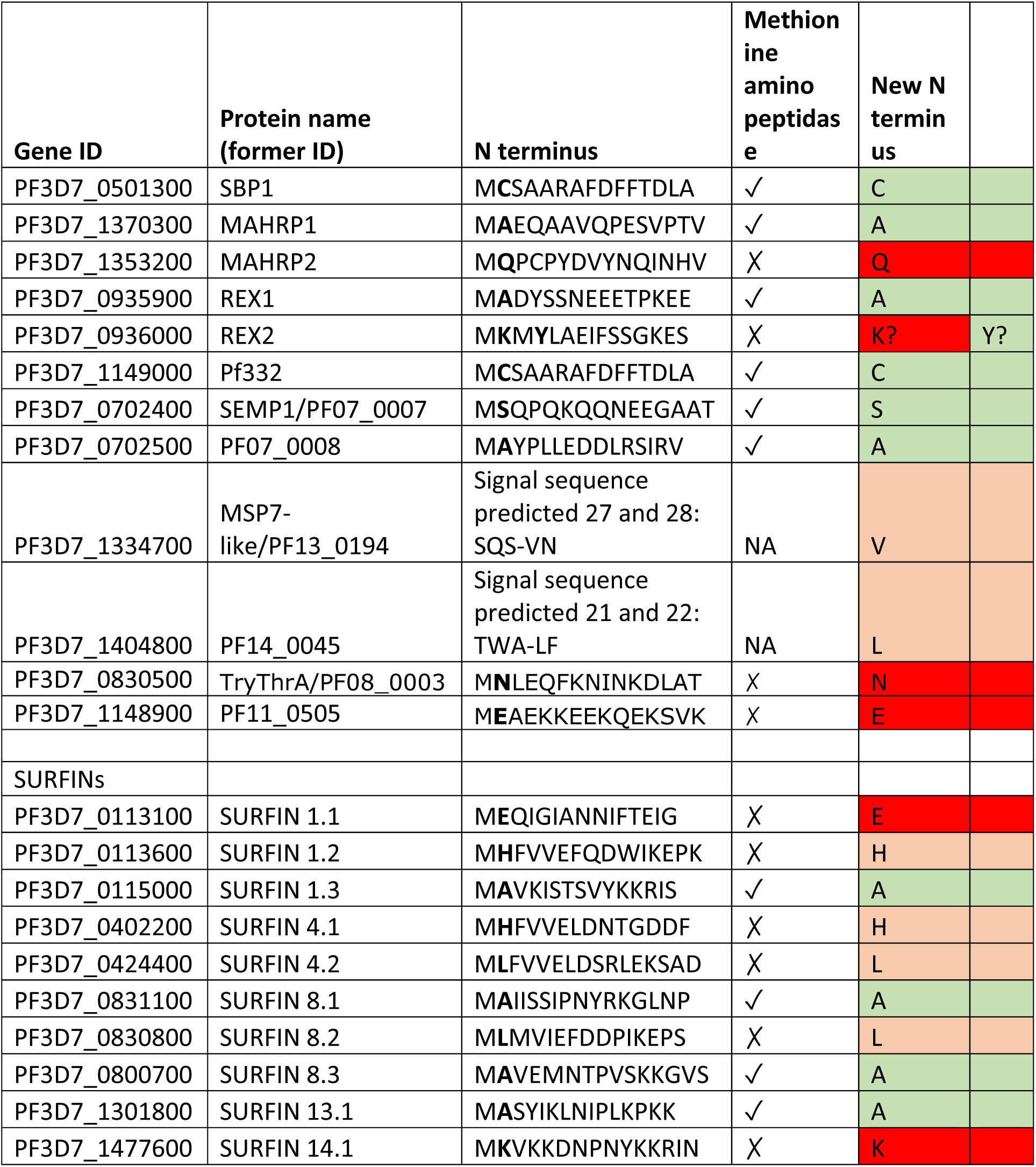
N termini of known PNEPs and the SURFIN family members. . Indicated are the N termini of the proteins, whether the start methionine is likely to be removed and the expected N-terminal residue after processing by methionine aminopeptidase. Color coding from Table 1 for N-terminal amino acids of PEXEL proteins: green-common, confirmed export; orange-uncommon, confirmed export; red-none confirmed, unlikely; grey-not observed. NA-not applicable.

Considering this bias towards certain N-terminal residues in exported proteins, we next determined whether these residues are also common at the N termini of proteins that are not exported. Interestingly, analysis of the predicted N termini of 73 PV, PVM and parasite surface proteins (which are processed by signal peptidase rather than PM V) revealed that frequently the predicted N-terminal residue is a large, charged residue, whereas small, polar residues are less common (Table 3, Supp Table 5) – over 43% (30/70) of these proteins have a predicted N-terminal residue that is in the ‘None confirmed – unlikely’ category of PEXEL proteins (i.e., an E, F, K or R residue) and second largest category (22/70) is ‘Uncommon-confirmed’ (i.e., a G, H, L, N or V residue). Only 20% (14/70) of the proteins contained a predicted N-terminal residue that is common in the fourth position of exported PEXEL proteins (‘Common-confirmed’ (i.e., an A, C, S, T or Y residue)). Included among these are the exported proteins Clag3.1 (S) and MSRP6 (S) (Beck et al., 2014; Nguitragool et al., 2014; Soares et al., 2021) (although there are different reports about the requirement of the PTEX for the export of Clag3 and the RhopH proteins (Beck et al., 2014; Schureck et al., 2021)). Note that in three proteins no signal sequence was predicted by SignalP. Interestingly, although the residue in the fourth position of the PEXEL is not conserved among PV6 orthologues in other *Plasmodium* species, nearly all contain a large, charged residue at this position (Supp Table 2); only in the *Plasmodium gallinaceum* orthologue were we unable to find a canonical PEXEL sequence.

**Table 3.**
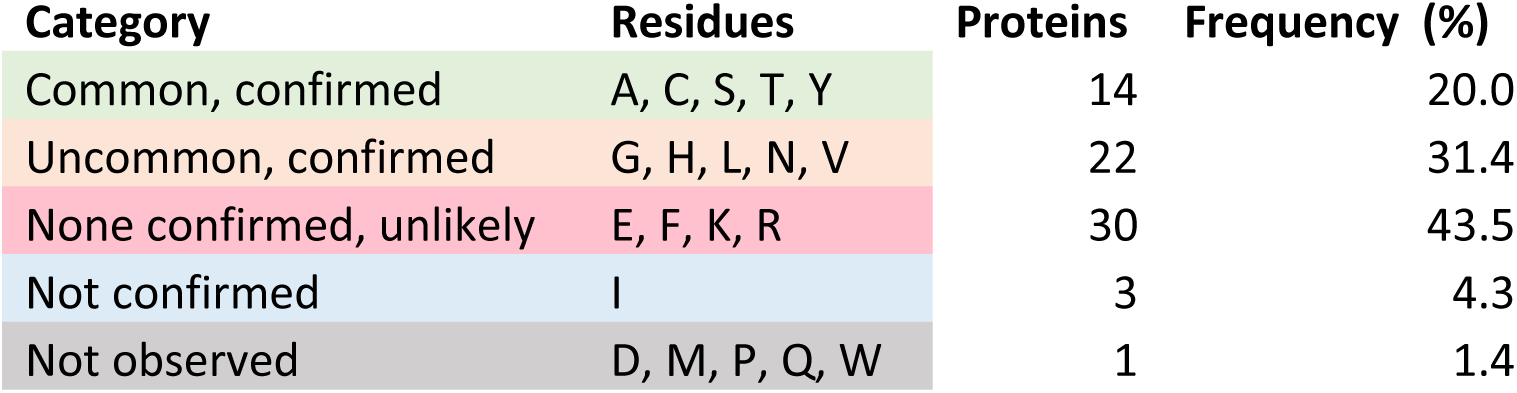
Predicted N-terminal amino acids of secreted apical organelle proteins, parasite surface proteins, parasitophorous vacuole proteins and parasitophorous vacuole membrane proteins. For full list of proteins, please see Supplementary Table 5.

These results are consistent with previous findings that the N-terminal residue of proteins reaching the PV and exposed to the PTEX likely contributes whether the protein will be exported. It also shows that the three pathways that the parasite uses to translocate and process proteins into the ER for transport to the PV – signal peptidase-mediated processing, PM V-mediated processing and post-translational transfer into the ER (in the case of PNEPs) – tend to expose different residues at new N terminus, potentially explaining how the different pathways promote export or retention in the PV.

### At least one additional Plasmepsin V target is essential for parasite development immediately after invasion

A previous study revealed that treatment of malaria parasites with a PM V inhibitor late in the erythrocytic cycle leads to a phenotype very early in the next erythrocytic cycle, much earlier than the ring-trophozoite stage arrest detected in mutants deficient in protein export (Beck et al., 2014; Elsworth et al., 2014; Charnaud et al., 2018; Polino et al., 2020). As AMA1-HA_3_-PV6 relies on signal peptidase rather than PM V for its processing, it afforded the opportunity to determine whether the ring-stage phenotype detected after treatment with PM V inhibitors is the result of the lack of processing of PV6 or whether additional PEXEL proteins are required for early ring-stage development. We treated synchronized wildtype 3D7 parasites and parasites producing AMA1-HA_3_-PV6 with the PM V inhibitor 601 during the schizont stage (Fig 5A) and inhibited egress with ML10 to control the timing of invasion (Baker et al., 2017; Ressurreição et al., 2020). Development of the schizonts did not appear to be affected in either parasite strain by the presence of 601, as judged by observation of Giemsa-stained parasites (Fig 5A). We then removed the ML10 and 601 to allow egress and invasion. In the case of the parasites treated only with ML10, we added 601 to half of the culture to determine whether 601 affects invading parasites and early rings. Parasite treated only with ML10 and parasites exposed to 601 at the time of removal of ML10 (approximately 20 minutes before egress and subsequent invasion (Ressurreição et al., 2020)) developed into normal ring-stage parasites. In contrast, parasites that had been treated with 601 during schizogony (but invaded in the absence of the inhibitor) showed a very severe developmental defect. Rather than forming the characteristic ring shapes, parasites of both strains were detected as very compact spots (Fig 5A). Measurement of the size of the parasites showed that there was a significant size difference between rings formed by parasites treated with 601 as schizonts (Fig 5A, panel iii and Fig 5B) and untreated parasites and parasites treated during egress (Fig 5A, panels i and ii, respectively, and Fig 5B). No difference in the effect of the inhibitor was detected between wildtype parasites and parasites that encode the AMA1-HA_3_-PV6 protein (Fig 5B), indicating that even though the AMA1-HA_3_-PV6 parasites produce a version of PV6 that does not rely on PM V for its processing (Fig 3D), the parasites were sensitive to inhibition of PM V. Interestingly, the phenotype of wildtype parasites arrested by 601 did not phenocopy the phenotype of the *pv6* mutant (Fig 1A); the inhibitor-treated parasites formed small, compact parasites (Fig 5A, panels iii and vi), as also previously reported (Polino et al., 2020), whereas parasites lacking PV6 formed slightly more expanded rings, often with a translucent center (Fig 1A). Furthermore, the phenotype displayed by the treated parasites did not phenocopy that of parasites in which the related proteases Plasmepsin IX or Plasmepsin X was inactivated; lack of activity of those proteases causes an egress defect (Plasmepsin X) or invasion defect (Plasmepsin IX) and thereby prevents invasion (Nasamu et al., 2017; Pino et al., 2017; Favuzza et al., 2020). Hence, although we cannot completely rule out that the inhibitor has alternative targets, the phenotype observed is not consistent with inhibition of Plasmepsin IX and Plasmepsin X.

**Figure 5.**
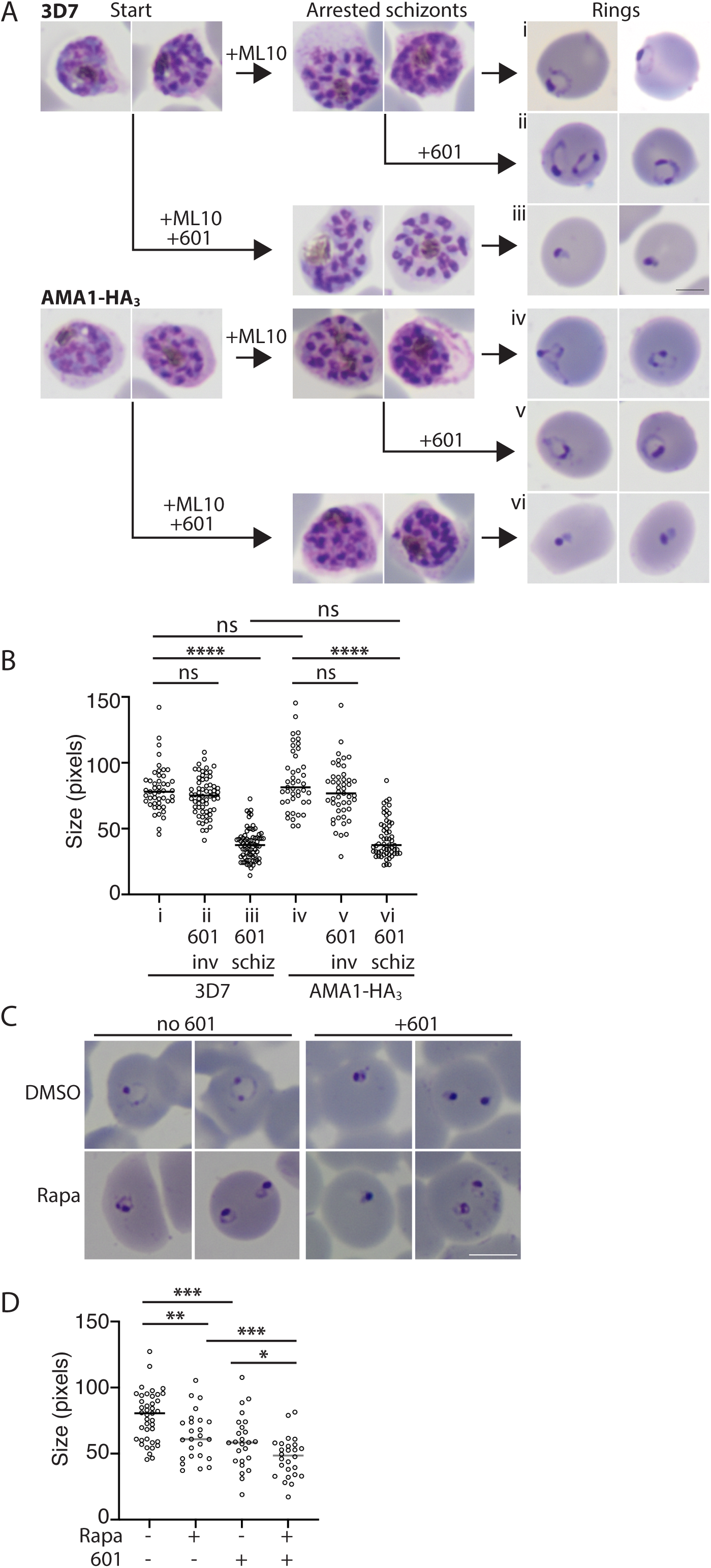
Developmental arrest induced by Plasmepsin V inhibitors after invasion. A) Synchronized cultures of 3D7 and AMA1-HA_3_-PV6 parasites were treated with ML10 when the parasites were in the early schizont stage and half of the culture was simultaneously treated with the PM V inhibitor WEHI-1252601 (601) (Start). ML10 and 601 were removed when most schizonts appeared arrested (Arrested schizonts). The culture treated only with ML10 was split into two and one half was treated with 601. Ring formation was determined one hour after removal of ML10 (Rings). Scale bar in right-hand side of panel iii represents 2.5 µm. B) Quantitation of parasite diameter size in samples i-vi in panel A. Indicated is the time the PM V inhibitor 601 was added; inv-after removal of ML10 from arrested schizonts; schiz-at start in Panel A (and removed with ML10). Data are based on the measurement of at least 40 rings. Mann–Whitney U test was performed for statistical analysis; ns: not significant, ****p < 0.0001. C) Treatment of parasites containing a floxed *pv6* locus with rapamycin and 601, as indicated. Note the more compact shape of the parasites treated with 601 compared to the rapamycin-treated parasites. Scale bar represents 5 µm. D) Quantitation of parasites sizes from experiment shown in panel C. Data are based on the measurement of at least 20 rings. Mann–Whitney U test was performed for statistical analysis: ns: not significant, *p < 0.05, **p < 0.01 ***p < 0.001, ****p < 0.0001.

To compare the phenotypes of parasites lacking PV6 and parasites in which PM V is inhibited directly, we repeated the experiment, treating PV6-diCre parasites carrying the floxed version of *pv6*, with DMSO and rapamycin and with 601. This showed once again that parasites lacking PV6 form small, round rings that often have a small translucent center. In contrast, parasites treated with 601 showed a more severe phenotype, with more compact parasites that in many cases did not have a clear center (Fig 5C). Quantitation of the size of the parasites further underscored the difference in size between the wild type, the mutant and the inhibitor-treated parasites (Fig 5D).

Together the findings that parasites expressing AMA1-HA_3_-PV6 remain sensitive to a PM V inhibitor, with a similar phenotype to that of wildtype parasites after invasion, and the difference in the morphology of parasites treated with the PM V inhibitor and parasites lacking PV6 indicate that at least one additional non-exported PM V substrate functions very soon after invasion.

## DISCUSSION

In this study we show that despite being cleaved by PM V, the *P. falciparum* phospholipid transfer protein PV6 is retained and functions in the PV. This is the first demonstration that a native PM V substrate in malaria parasites is not exported into the host cell and shows that cleavage by PM V does not always per se target a protein for export. Furthermore, we show that the fourth residue of the PEXEL in PV6, a K, which becomes the N-terminal residue after cleavage by PM V, determines the localization of the protein; when this residue is changed to an A – a residue found in the fourth position of several exported proteins, including KAHRP (Boddey et al., 2009) – the protein is exported. Previous studies have shown that altering this position in exported proteins can severely decrease the fraction of the protein that is exported without affecting cleavage by PM V (Grüring et al., 2012; Tarr et al., 2013). However, PV6 is the first native PM V substrate identified that has a demonstrated essential function in the PV. This finding indicates that processing of a protein by PM V does not inevitably result in protein export and that processing by PM V and recognition of proteins for export seem to be two separate events that are unlikely to be mechanistically coupled.

Our results furthermore provide additional support for the finding that the main determinant of protein export is the N-terminal residue (Grüring et al., 2012; Tarr et al., 2013), as long as this is followed by a flexible region of approximately 10 amino acids and the protein can be unfolded by HSP101 (Knuepfer et al., 2005a; Gehde et al., 2009; Matthews et al., 2019), although additional residues following the N terminus also influence protein export (Heiber et al., 2013; Tarr et al., 2013; Chitama et al., 2019). PEXEL proteins for which export has been experimentally verified most commonly contain a small, polar residue at the fourth position of the PEXEL (Table 1) (Hiller et al., 2004; Marti et al., 2004). Furthermore, the PNEPs analyzed in this study contain in many cases a small, polar residue in the second position – these residues are also favored by methionine aminopeptidase for removal of the N-terminal M residue and will therefore likely become the N terminus after processing by the peptidase. In contrast, processing of parasite surface proteins, PV proteins, PVM proteins and apical organelle proteins by signal peptidase is predicted to expose in many cases a large, charged residue at the newly formed N terminus – residues that are not found in known exported proteins. One notable exception is the exported protein MSRP6 (Beck et al., 2014; Soares et al., 2021), in which an S is predicted to become the N-terminal residue after processing by signal peptidase. Based on this, we propose a model in which the N-terminal region of a protein with a small, polar residue at the extreme N terminus is recognized by the PTEX (potentially HSP101 or an accessory factor) and thereby targets these for export, regardless of the process by which the N terminus has been generated. Proteins that feature a large, charged residue at the N terminus, such as many surface proteins, PV proteins and PVM proteins that are processed by signal peptidase, are not recognized and hence are not exported. As parasite surface proteins, PV proteins and PVM proteins that are cleaved by signal peptidase often contain a large, charged residue at the N terminus after cleavage, they would not be recognized by the export machinery, thereby preventing improper removal of proteins from either the surface of the parasite or the PV or clogging the PTEX with parasite surface proteins. The fate of proteins that feature at their N terminus an amino acid that is present in exported and in surface proteins may be influenced by the residues that follow the N-terminal residue, as these also play an important role in determining the localization of parasite proteins (Grüring et al., 2012; Tarr et al., 2013; Chitama et al., 2019).

Together, our results uncouple processing by PM V from protein export and indicate that the role of PM V in *Plasmodium* parasites is broader, and likely more ancient, than processing exported proteins, as has been postulated previously (Nasamu et al., 2020). Its role therefore may be more similar to that of its *Toxoplasma gondii* orthologue, the aspartic protease ASP5, than previously thought. One notable difference between the two organisms is that nearly all ASP5 substrates function in the PV and the PVM (Hsiao et al., 2013; Coffey et al., 2018), whereas all PM V substrates were thought to be exported. Our findings that a PV protein is cleaved by PM V and that there is at least one other PM V target that does not function in the cytosol of the host erythrocyte may indicate that in *Plasmodium* parasites, PM V also functions to process PV and PVM proteins. As orthologues of PM V are conserved among many Apicomplexan parasites, we postulate that PM V evolved in Apicomplexan parasites to cleave PV or PVM proteins and that the role of PM V in processing exported proteins evolved subsequently in *Plasmodium* parasites. It is of note that conserved PEXEL proteins (including orthologues of PV6) generally contain a large, charged residue at position 4 of the PEXEL (van Ooij et al., 2008). It is possible that, contrary to what has been reported previously (van Ooij et al., 2008), most, if not all, conserved PEXEL proteins function as non-exported proteins, indicating that their functions may have arisen prior to the development of the export machinery in malaria parasites. Why the parasite has evolved to use two proteases, PM V and signal peptidase, to process proteins that enter the secretory pathway remains to be discovered.

## ACKNOWLEDGMENTS

We thank Justin Boddey and Brad Sleebs for sharing the PM V inhibitors, Eizo Takashima for sharing the anti-PV1 antiserum, Paul Gilson for sharing the rabbit anti-EXP2 antiserum and Mike Blackman for sharing antibodies against AMA1, EXP1, RAP1 and RON4. The anti-EXP2 monoclonal antibody 7.7 was obtained from the European Malaria Reagent Repository, the anti-RESA monoclonal antibody 28/2 was obtained from the antibody facility at the Walter and Eliza Hall Institute of Medical Research. We thank Angelika Gründling for her assistance in constructing plasmids pBLD706 and pBLD708. The authors acknowledge the facilities and the scientific and technical assistance of the LSHTM Wolfson Cell Biology Facility, with specific thanks to Liz McCarthy and Netanya Bernitz. This work was supported by a Medical Research Council Career Development Award to CvO (MR/R008485/1).

## SUPPLEMENTARY FIGURE LEGENDS

Supplementary Figure 1. **Excision PCR of parasites with floxed *pfa0210c* locus and costaining of parasites with antibodies against dense granule markers markers**. A). Excision PCR of the parasite line containing the floxed *pfa0210c* locus used in Figure 1, showing the *pfa0210* locus after parasites have been treated with DMSO (D) or rapamycin (R). See Hill *et al*. for outline of PCR strategy (Hill et al., 2016). B) Immunofluorescence assay of a lysed late-stage schizont (top) and free merozoite (bottom) with antibodies against PV6 (PFA0210c; green) and the dense granule marker EXP2 (red). The DNA was stained with Hoechst 33342 and is shown in blue. Scale bar top images: 5 µm; scale bar bottom images: 1 µm.

Supplementary Figure 2. **Immunofluorescence assays of erythrocytes containing recently invaded parasites.** A) Maximum projection view of infected erythrocytes shown in Figure 2B, bottom row, showing that both erythrocytes that display anti-RESA staining are infected. B) Co-staining of infected erythrocytes with anti-PV6 and anti-EXP2 antibodies after invasion. Note that although the proteins do not overlap to a large degree, they appear to be present in the same compartment.

Supplementary Figure 3. **Genetic modification of the *pfs47* locus and localization of PV6 K64A mutant protein.** A) overview of the integration strategy using Cas9-mediated gene modification of the *pfs47* locus with the *pv6* gene carrying the K64A mutation and PCR analysis of integration of parasites obtained. Indicated above are listed the primer pairs used in the PCR and underneath are the expected sizes of the PCR products (see also Primers Supp Table 1) B) Immunofluorescence assays of erythrocytes infected with wildtype parasites or parasites infected with parasites expressing an additional copy of PV6 containing a K64A mutation, as indicated. Cells were stained with antibodies against PV6 to visualize PV6 and either PV1 or EXP1 to visualize the PV or PVM, respectively. C) PCR analysis of the *pv6* locus of parasites carrying the K64A mutant at the *pfs47* locus after treatment with DMSO (D) or rapamycin (R). For outline of PCR scheme, please see Hill et al. (Hill et al., 2016)

